# *Piper longum*: A review of its phytochemicals and their network pharmacological evaluation

**DOI:** 10.1101/169763

**Authors:** Neha Choudhary, Vikram Singh

## Abstract

*Piper longum* L. (*P. longum*, also called as long pepper) is one of the common culinary herb and has been extensively used as an important constituent of various indigenous medicines, specifically in traditional Indian medicinal system known as Ayurveda. Towards obtaining a global regulatory framework of *P. longum*’s constituents, in this work we first reviewed phytochemicals present in this herb and then studied their pharmacological and medicinal features using network pharmacology approach. We developed high-confidence level tripartite networks consisting of phytochemicals – protein targets – disease association and explain the role of its phytochemicals to various chronic diseases. 7 drug-like phytochemicals in this herb were found as the potential regulators of 5 FDA approved drug targets; and 28 novel drug targets were also reported. 105 phytochemicals were linked with immunomodulatory potency by pathway level mapping in human metabolic network. A sub-network of human PPI regulated by its phytochemicals was derived and various modules in this sub-network were successfully associated with specific diseases.

**Graphical abstract:** 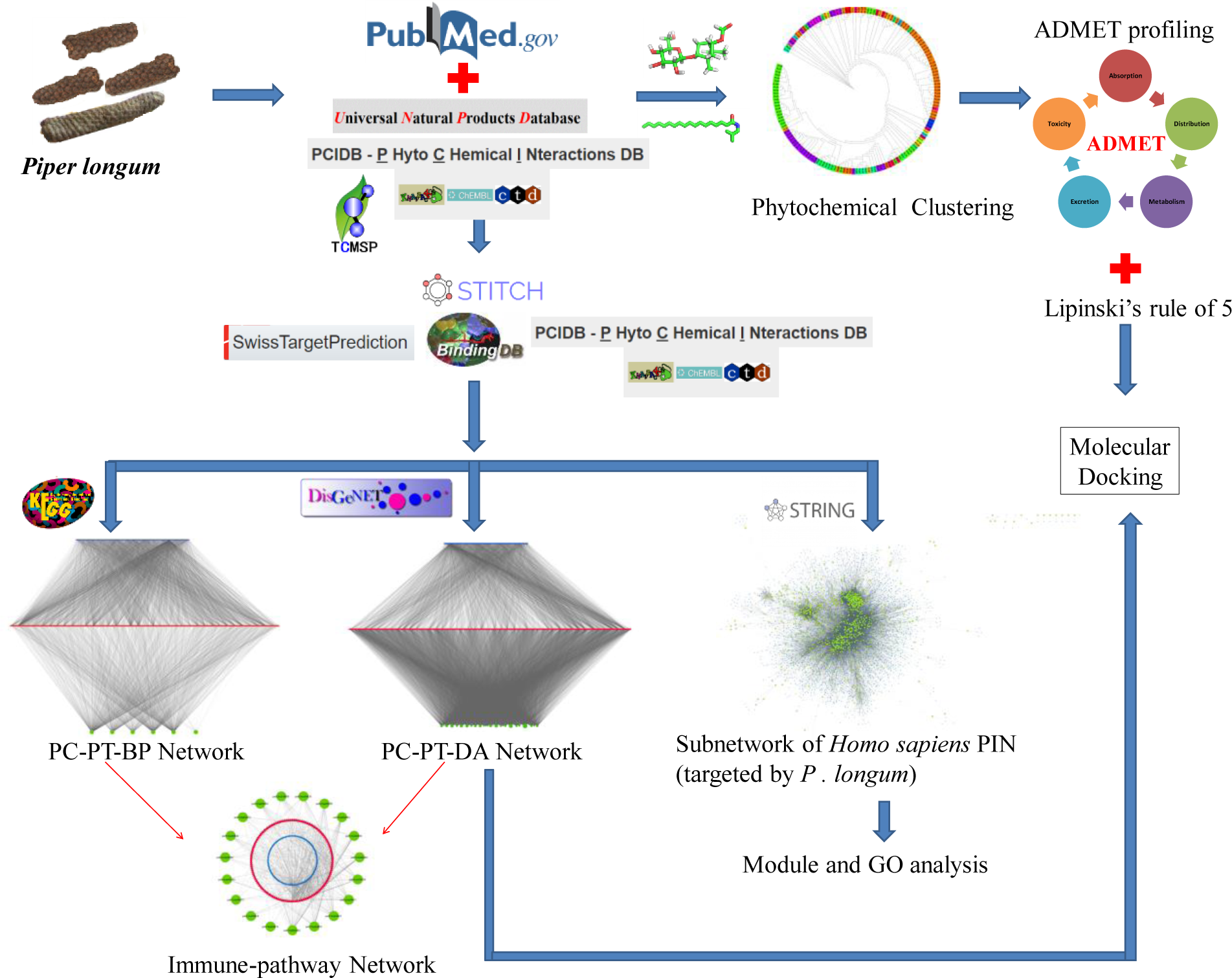

**Abbreviations:**

*P. longum*: *Piper longum* L.
PC: Phytochemical
PT: Protein target
BP: Biological pathways
DA: Disease asscociation
PCt: Number of protein targets corresponding to a particular phytochemical
Tt: Total number of protein targets of *P. longum*
ADMET: Absorption, Distribution, Metabolism, Excretion and Toxicity.

## 1. Introduction

Healing with medicinal plants is an ancient idea. Secondary metabolites of various plants have been traditionally utilized for the betterment of human-health. Plants belonging to genus Piper are amongst the important medicinal plants used in various systems of medicine. More than 1000 species belongs to this genus and *P. longum* is one of the most well-known species of it, including *Piper nigrum* and *Piper bettle*. *P. longum* forms an active constituent of the widely used Ayurvedic poly-herbal formulation ―Trikatu‖ (Johri et al., 1992). The wide spread use of this herb in different formulations as documented in ancient Ayurvedic manuscripts such as Caraka samhita (Das and Sharma, 2002), Susruta samhita (Srikantha Murthi, 2012), Vagbhata‘s astanga hrdayam (Srikantha Murthi, 2000) etc. suggests its importance in traditional Indian medicinal system.

*P. longum* is an indigenously growing plant in India and is also cultivated in the tropical and subtropical regions of Asia and Pacific islands (Tripathi et al., 1999). It is usually cultivated for its fruit, which is dried and used as spice. The plant grows into a shrub with large woody roots, numerous creeping and jointed stems that are thickened at the nodes. The leaves are without stipules and spreading in nature. Fruits are small and oval shaped berries, grown as spike. Ones matured, the spikes are collected. The dried form of spikes makes ―pippali‖ while its root radix is known as ―pippalimula‖. The dietary piperine is known for its bioavailability and digestive enhancing properties. *In vitro* studies have shown the role of piperine in relieving oxidative stress by quenching free radicals and reactive oxygen species. It is known to act as an antimutagenic and antitumor agent (Srinivasan, 2007). Anti-diarrheic and anti-dysenteric properties of this spice enhance its medicinal value (Reddy et al., 2001). The pharmacological properties of this plant include anticancer, antioxidant, anti-inflammatory, hepatoprotective, immunomodulatory, antimicrobial, anti-platelet, anti-hyperlipidemic, analgesic, antidepressant, anti-amoebic, anti-obesitic, radio-protective, cardio-protective and anti-fungal activities (Kumar et al., 2011; Sunil and Kuttan, 2004; Sharma et al., 2014; Jalalpure et al., 2003). Methanolic extract of this fruit has been reported to be involved in memory repairment and improving memory performance as shown by *in vitro* model (Hritcu et al., 2014). Clinical studies have revealed the efficacy of this plant in treatment of bronchial asthma in children (Clark et al., 2010; Dahanukar et al., 1984). Antidiabetic activity of the roots has also been reported. It is widely used as an important constituent in various Ayurvedic medicines to cure diseases like leprosy and tuberculosis and also used in the treatment of cough, dysponea, cardiac and spleen disorders, chronic-fever, gout, rheumatic pain etc. (Khushbu et al., 2011).

In recent years, the advancement in chemistry, pharmacology and systems biology has created a new paradigm for the drug discovery that is known as Network pharmacology (Hopkins, 2008). The integration of traditional knowledge of medicine with recent *in silico* approaches has led to the identification of novel natural drug compounds. The approach has recently gathered much attention by the research community as network pharmacology based studies have been widely used to explore the medicinal activities of herbs like Withania somnifera (Chandran et al., 2017) and formulae like QiShenYiQi (Li et al., 2014(a)), Ge-gen-qin-lian decoction (Li et al., 2014(b)) etc. to understand their molecular level effect in the treatment of syndrome or diseases.

In the present work, we have reviewed the phytochemicals of *P. longum* as reported in literature and public databases and studied the pharmaceutically relevant features of these phytochemicals. The therapeutic relevance of these compounds was inferred through the network analysis of phytochemicals with their protein targets. The therapeutic activity of the phytochemicals was correlated with the number of proteins that the molecules may target. Further, the pharmacological action of these metabolites at biological level was explored and the potential metabolic and cellular pathways in which the target proteins are involved have been identified. Disease association network was constructed to interpret the relationship between the potential drug candidates in the human system. Finally, a subnetwork of human protein-protein interaction (PPI) network that is potentially regulated by *P. longum* was analyzed to identify functional modules present therein.

## 2. Materials and methods

### 2.1. Data collection

A dataset of phytochemicals present in *P. longum* was developed using extensive literature survey and mining of public database resources. Relevant research articles from pubMed-NCBI (https://www.ncbi.nlm.nih.gov/pubmed/) were selected and manually scrutinized. Three databases UNPD (Universal Natural Products Database) (pkuxxj.pku.edu.cn/UNPD), TCMSP (Traditional Chinese Medicine Systems Pharmacology) (Ru et al., 2014) and PCIDB (PhytoChemical Interactions DB) (http://www.genome.jp/db/pcidb) were screened for the potentially active phytochemical present in *P. longum*. Chemical information of these phytochemicals was compiled from PubChem database (Bolton et al., 2008) and ChEMBL (Gaulton et al., 2011). Structural data of phytochemical not available in PubChem and ChEMBL was derived using PubChem Sketcher V2.4 (Ihlenfeldt et al., 2009). The compiled dataset was filtered to remove duplicate entries. Human proteins targeted by the phytochemicals were predicted from STITCH 5.0 (Search Tool for Interactions of Chemicals and proteins) (Szklarczyk et al., 2015), BindingDB (Liu et al., 2006), SwissTargetPrediction (Gfeller et al., 2014) and PCIDB.

The protein-chemical interaction reported in STITCH comes from the manually curated datasets including DrugBank (Wishart et al., 2006), GLIDA (Okuno et al., 2006), Matador (Günther et al., 2007), TTD (Therapeutic target database) (Chen et al., 2002) etc. Information from several metabolic pathway databases and experimentally validated interactions from ChEMBL (Gaulton et al., 2011), PDB (Kouranov et al., 2006) and others sources provides a full picture of the all available data. In order to access the high confidence targets, the interactions with combined score of ≥0.4 were taken into account. BindingDB has the basic working principle that similar compounds tend to bind same proteins, and is a web accessible database containing binding affinities of 20,000 (approx.) protein-ligand complexes. We screened the phytochemical-protein interactions having similarity search value ≥0.85. SwissTargetPrediction uses a combination of 2D and 3D similarity measures to identify the target proteins that can bind with a ligand showing the highest similarity with a library of 280,000 compounds. Top 15 protein targets for each phytochemical were selected from this resource. PCIDB returns a list of active compounds of the query herb and also enlists the possible genes involved in the interactions. We included all the phytochemical-protein interactions provided by PCIDB. Uniprot IDs of all the protein targets identified from the above mentioned four resources were used for network construction.

The biological pathways association of the identified protein targets was retrieved from KEGG database (Kyoto Encyclopedia of Genes and Genomes) (Kanehisa et al., 2000). A comprehensive platform of gene-disease association, called DisGeNETv4.0 (Piñero et al., 2017) was used to find the disease information in which the protein targets may be involved. DrugBank database (Wishart et al., 2006) was also mined to find-out the known target proteins.

### 2.2. Compounds classification and clustering

An automated and rapid chemical classification method ―ClassyFire‖ (Feunang et al., 2016) was used to assign chemical class to the phytochemicals. The query is mapped to the various classes based on its features that are calculated using superstructure-search operations and other properties. Clustering toolbox from ChemMine tools (Backman et al., 2011) was used for clustering the phytochemicals. The hierarchical clustering algorithm was opted, that forms a hierarchy of clusters based on pairwise compound similarities using the atom-pair descriptors and Tanimoto coefficient.

### 2.3. Network construction and analysis

To investigate the pharmacological actions of the phytochemicals, various networks showing the interactions among phytochemical compounds (PC), protein targets (PT), biochemical pathways (BP) and associated diseases (AD) were constructed and analyzed using Cytoscape v3 (Shannon et al., 2003).

### 2.4. Modularity analysis of human PPI subnetwork

STRING v10.5 (Szklarczyk et al., 2014) was used to identify first degree interactors of all the target proteins. Only high confidence interactions (score >=0.9) were included to construct the PPI and duplicate edges were removed. MCL (Markov Cluster) algorithm (Enright et al., 2002) was used to cluster and organize the proteins in various modules. This algorithm detects cluster structure using mathematical bootstrapping procedure and has been shown that it is efficient in identification of modules in PPI networks (Brohee et al., 2006). GO based functional enrichment of these clusters was carried out using BINGO (Maere et al., 2005).

### 2.5. Drug-likeliness prediction and molecular docking studies

Pharmacokinetic and toxicity properties of the compounds were studied using pKCSM which uses graph based structure signature method to predict a range of ADMET(Absorption, Distribution, Metabolism, Excretion, Toxicity) properties (Pires et al., 2015).

AutoDock 4.2 (Morris et al., 2014) was used to carry out the computational docking studies, using Lamarckian Genetic algorithm. The best binding orientation of the ligand within the protein cavity was estimated using binding energy values.

## 3. Results and discussion

### 3.1. Identification of phytochemical compounds

Phytochemical data of *P. longum* were obtained from extensive literature survey and mining natural product databases. In total 159 phytochemicals were identified and all the phytochemicals were assigned a unique ID. Details of all the phytochemicals i.e. their unique ID, name and their reference are presented in Table 1.

**Table 1.** List of phytochemicals identified in *P. longum*.

| S. No. | Phytochemical ID | Phytochemical name | Pubchem/ ChEMBL ID | Reference |
| --- | --- | --- | --- | --- |
| 1 | PL1 | 3β, 4α-dihydroxy-1-(3-phenylpropanoyl)-piperidine-2-one | N/A | Jiang et al., 2013 (a) |
| 2 | PL2 | (2E, 4E, 14Z)-6-hydroxyl-N-isobutyleicosa-2,4,14-trienamide | N/A | Jiang et al., 2013 (a) |
| 3 | PL3 | Coumapherine | 10131321 | Liu et al., 2009 |
| 4 | PL4 | N-5-(4-hydroxy-3-methoxyphenyl)-2E-pentenoyl piperidine | N/A | Liu et al., 2009 |
| 5 | PL5 | Piperolactam A | 3081016 | Liu et al., 2009 |
| 6 | PL6 | 1-[1-oxo-5 (3,4-methylenedioxyphenyl) -2E,4E-pentadienyl] -piperidine | N/A | Liu et al., 2009 |
| 7 | PL7 | (R)-(-) –turmerone | 558221 | Liu et al., 2009 |
| 8 | PL8 | Octahydro-4-hydroxy-3α-methyl-7-methylene-α-(1-methylethyl)-1H-indene-1-methanol | N/A | Liu et al., 2009 |
| 9 | PL9 | (+) -aphanamol I | 11031884 | Liu et al., 2009 |
| 10 | PL10 | Bisdemethoxycurcumin | 5315472 | Liu et al., 2009 |
| 11 | PL11 | Demethoxycurcumin | 5469424 | Liu et al., 2009 |
| 12 | PL12 | Longumosides A | N/A | Jiang et al., 2013 (b) |
| 13 | PL13 | Longumosides B | 71579641 | Jiang et al., 2013 (b) |
| 14 | PL14 | Erythro-1-[1-oxo-9(3,4-methylenedioxyphenyl)-8,9-dihydroxy-2E-nonenyl]-piperidine | N/A | Jiang et al., 2013 (b) |
| 15 | PL15 | Threo-1-[1-oxo-9(3,4-methylenedioxyphenyl)-8,9-dihydroxy-2E-nonenyl]-piperidine | N/A | Jiang et al., 2013 (b) |
| 16 | PL16 | 3β,4α-dihydroxy-2-piperidinone | N/A | Jiang et al., 2013 (b) |
| 17 | PL17 | 5,6-dihydro-2(1H)-pyridinone | N/A | Jiang et al., 2013 (b) |
| 18 | PL18 | Piperlongumide (1) [N-isobutyl-19-(3',4'-methylenedioxyphenyl)-2E,4E nonadecadienamide] | N/A | Sahi et al., 2013 |
| 19 | PL19 | 1-(3,4-methylenedioxyphenyl)-1E tetradecene | N/A | Sahi et al., 2013 |
| 20 | PL20 | Piperlongimin A [2E-N-isobutyl-hexadecenamide] | N/A | Sahi et al., 2013 |
| 21 | PL21 | 2E,4E-N-isobutyl-octadecenamide | N/A | Sahi et al., 2013 |
| 22 | PL22 | Piperlongimin B [2E-octadecenoylpiperidine] | N/A | Sahi et al., 2013 |
| 23 | PL23 | 2E,4E-N-isobutyl-dodecenamide | N/A | Sahi et al., 2013 |
| 24 | PL24 | 2E,4E,12E,13-(3,4-methylenedioxyphenyl)-trideca-trienoic acid isobutyl amide | N/A | Sahi et al., 2013 |
| 25 | PL25 | Piperine | 638024 | Sahi et al., 2013 |
| 26 | PL26 | Pellitorine | 5318516 | Li et al., 2013 |
| 27 | PL27 | N-[(2E,4E)-Decadienoyl]-piperidine | 11118018 | Li et al., 2013 |
| 28 | PL28 | N-Isobutyl-2E,4E-undecadienamide | 20157325 | Li et al., 2013 |
| 29 | PL29 | Piperlonguminine | 5320621 | Li et al., 2013 |
| 30 | PL30 | Piperanine | 5320618 | Li et al., 2013 |
| 31 | PL31 | N-[(2E,4E)-Tetradecadienoyl]piperidine | 11130083 | Li et al., 2013 |
| 32 | PL32 | N-Isobutyl-2E,4E-hexadecadienamide | 6442402 | Li et al., 2013 |
| 33 | PL33 | Pipercollosine | 5372201 | Li et al., 2013 |
| 34 | PL34 | (2E,4E,12Z)-N-Isobutyl-octadeca-2,4,12-trienamide | N/A | Li et al., 2013 |
| 35 | PL35 | N-Isobutyl-2E,E-octadecadienamide | 9974234 | Li et al., 2013 |
| 36 | PL36 | Dehydropipernonaline | 6439947 | Li et al., 2013 |
| 37 | PL37 | Pipernonatine | 9974595 | Li et al., 2013 |
| 38 | PL38 | (E)-9-(Benzo[d][1,3]dioxol-5-yl)-1-(piperidin-1-yl)non-2-en-1-one | N/A | Li et al., 2013 |
| 39 | PL39 | 1-(2E,4E,12E)-Octadecatrinoylpiperidine | N/A | Li et al., 2013 |
| 40 | PL40 | Retrofractamide B | 5372162 | Li et al., 2013 |
| 41 | PL41 | (2E,4E,14Z)-N-Isobutyleicosa-2,4,14-trienamide | N/A | Li et al., 2013 |
| 42 | PL42 | N-isobutyl-2E,4E-decyldecadienamide | N/A | Li et al., 2013 |
| 43 | PL43 | (2E,4E,10E)-N-11-(3,4-Methylenedioxyphenylhmdecatrienoylpiperidine | N/A | Li et al., 2013 |
| 44 | PL44 | 1-[(2E,4E,14Z)-1-Oxo-2,4,14-eicosatrienyl]-piperidine | N/A | Li et al., 2013 |
| 45 | PL45 | Guineensine | 6442405 | Li et al., 2013 |
| 46 | PL46 | (2E,4E,14Z)-N-Isobutyl docosa-2,4,14-trienamide | N/A | Li et al., 2013 |
| 47 | PL47 | (2E,4E,12E)-13-(Benzo[d][1,3]dioxol-6-yl)-1-(piperidin-1-yl)trideca-2,4,12-trien-1-one | N/A | Li et al., 2013 |
| 48 | PL48 | (2E,4E,13E)-14-(Benzo[d][1,3]dioxol-6-yl)-N-isobutyltetradeca-2,4,13-trienamide | N/A | Li et al., 2013 |
| 49 | PL49 | Brachyamide B | 14162526 | Li et al., 2013 |
| 50 | PL50 | Dihydropiperlonguminine | 12682184 | Li et al., 2013 |
| 51 | PL51 | Piperdardine | 10086948 | Li et al., 2013 |
| 52 | PL52 | Retrofractamide A | 11012859 | Li et al., 2013 |
| 53 | PL53 | Piperchabamide D | 16041827 | Li et al., 2013 |
| 54 | PL54 | N-isobutyl-2E,4E-dodecadienamide | 6443006 | Li et al., 2013 |
| 55 | PL55 | Piperchabamide B | 44453655 | Li et al., 2013 |
| 56 | PL56 | 13-(1,3-Benzodioxol-5-yl)-N-(2-methylpropyl)-(2E,4E)-tridecadienamide | N/A | Li et al., 2013 |
| 57 | PL57 | Piperchabamide C | 44454018 | Li et al., 2013 |
| 58 | PL58 | 1-[(2E,4E)-1-oxo-2,4-hexadecadienyl]-piperidine | 10980124 | Li et al., 2013 |
| 59 | PL59 | 2,2-Dimethoxybutane (C6H14O2] | 137941 | Das et al., 2012 |
| 60 | PL60 | 2-Hydroxy myristic acid (C14H28O3] | 1563 | Das et al., 2012 |
| 61 | PL61 | β-Myrcene (C10H16] | 31253 | Das et al., 2012 |
| 62 | PL62 | N-methyl-1- octadecanamine (C19H41N] | 75539 | Das et al., 2012 |
| 63 | PL63 | Piperazine adipate (C10H20N2O4] | 8905 | Das et al., 2012 |
| 64 | PL64 | 2-Nonynoic acid (C9H14O2] | 61451 | Das et al., 2012 |
| 65 | PL65 | Dodecanal (CH3(CH2]10CHO] | 8194 | Das et al., 2012 |
| 66 | PL66 | 1,2-Benzenedicarboxylic acid, bis(2-ethylhexyl] ester (C24H38O4] | 8343 | Das et al., 2012 |
| 67 | PL67 | 2-Amino-4-hydroxypteridine-6-carboxylic acid (C7H5N5O3] | 70361 | Das et al., 2012 |
| 68 | PL68 | Piperlongumine | 637858 | Chatterjee et al., 1967 |
| 69 | PL69 | Hydrochloric acid (HCl) | 107 | Ru et al., 2014 |
| 70 | PL70 | Palmitic acid | 985 | Ru et al., 2014 |
| 71 | PL71 | 1,8-cineole | 2758 | Ru et al., 2014 |
| 72 | PL72 | Lawson | 6755 | Ru et al., 2014 |
| 73 | PL73 | Cis-Decahydronaphthalene | 7044 | Ru et al., 2014 |
| 74 | PL74 | Piperonylic acid | 7196 | Ru et al., 2014 |
| 75 | PL75 | Hypnon | 7410 | Ru et al., 2014 |
| 76 | PL76 | Moslene | 7461 | Ru et al., 2014 |
| 77 | PL77 | Cymol | 7463 | Ru et al., 2014 |
| 78 | PL78 | Methyl hydrocinnamate | 7643 | Ru et al., 2014 |
| 79 | PL79 | Hexahydropyridine (PIP) | 8082 | Ru et al., 2014 |
| 80 | PL80 | Pisol | 8193 | Ru et al., 2014 |
| 81 | PL81 | Piperonal | 8438 | Ru et al., 2014 |
| 82 | PL82 | Isobutylisovalerate | 11514 | Ru et al., 2014 |
| 83 | PL83 | Tridecane (TRD) | 12388 | Ru et al., 2014 |
| 84 | PL84 | Pentadecane (MYS) | 12391 | Ru et al., 2014 |
| 85 | PL85 | N-Heptadecane | 12398 | Ru et al., 2014 |
| 86 | PL86 | N-Nonadecane (UPL) | 12401 | Ru et al., 2014 |
| 87 | PL87 | Tridecylene | 17095 | Ru et al., 2014 |
| 88 | PL88 | Heptadecene | 23217 | Ru et al., 2014 |
| 89 | PL89 | Pentadecene | 25913 | Ru et al., 2014 |
| 90 | PL90 | Nonadecene | 29075 | Ru et al., 2014 |
| 91 | PL91 | tetradecadiene-1,13 | 30875 | Ru et al., 2014 |
| 92 | PL92 | Linalool (D) | 67179 | Ru et al., 2014 |
| 93 | PL93 | Cyclopentadecane | 67525 | Ru et al., 2014 |
| 94 | PL94 | Beta-Bisabolene | 68128 | Ru et al., 2014 |
| 95 | PL95 | Sesamol | 68289 | Ru et al., 2014 |
| 96 | PL96 | Sesamin | 72307 | Ru et al., 2014 |
| 97 | PL97 | p-Amino-o-cresol | 76081 | Ru et al., 2014 |
| 98 | PL98 | 2,4-Dimethoxytoluene | 96403 | Ru et al., 2014 |
| 99 | PL99 | D-Camphor (CAM) | 159055 | Ru et al., 2014 |
| 100 | PL100 | Cis-2-Decalone | 246289 | Ru et al., 2014 |
| 101 | PL101 | Piperitenone | 381152 | Ru et al., 2014 |
| 102 | PL102 | (-)-Nopinene | 440967 | Ru et al., 2014 |
| 103 | PL103 | (-)-Alpha-Pinene | 440968 | Ru et al., 2014 |
| 104 | PL104 | Isodiprene (CHEBI:7) | 443156 | Ru et al., 2014 |
| 105 | PL105 | (R)-linalool | 443158 | Ru et al., 2014 |
| 106 | PL106 | N-(2,5-dimethoxyphenyl)-4-methoxybenzamide | 532276 | Ru et al., 2014 |
| 107 | PL107 | Anethole | 637563 | Ru et al., 2014 |
| 108 | PL108 | Isoeugenol | 853433 | Ru et al., 2014 |
| 109 | PL109 | (3S)-3,7-dimethylocta-1,6-dien-3-yl] propanoate | 1616358 | Ru et al., 2014 |
| 110 | PL110 | ()-Terpinen-4-ol | 2724161 | Ru et al., 2014 |
| 111 | PL111 | Alpha-Farnesene | 5281516 | Ru et al., 2014 |
| 112 | PL112 | Farnesene | 5281517 | Ru et al., 2014 |
| 113 | PL113 | alpha-humulene | 5281520 | Ru et al., 2014 |
| 114 | PL114 | Isocaryophyllene | 5281522 | Ru et al., 2014 |
| 115 | PL115 | p-Ocimene | 5281553 | Ru et al., 2014 |
| 116 | PL116 | 8-Heptadecene | 5364555 | Ru et al., 2014 |
| 117 | PL117 | 9,17-OCTADECADIENAL (Z) | 5365667 | Ru et al., 2014 |
| 118 | PL118 | Cyclodecene, 1-methyl- | 5367581 | Ru et al., 2014 |
| 119 | PL119 | 1,4,7,-Cycloundecatriene, 1,5,9,9-tetramethyl-, Z,Z,Z,- | 5368784 | Ru et al., 2014 |
| 120 | PL120 | (+/-)-Isoborneol | 6321405 | Ru et al., 2014 |
| 121 | PL121 | (Z)-caryophyllene | 6429301 | Ru et al., 2014 |
| 122 | PL122 | -cis-.beta.-Elemene diastereomer | 6431152 | Ru et al., 2014 |
| 123 | PL123 | N-Isobutyl-2,4-icosadienamide | 6441067 | Ru et al., 2014 |
| 124 | PL124 | (E,E,E)-11-(1,3-Benzodioxol-5-yl)-N-(2-methylpropyl)-2,4,10-undecatrienenamide | 6453083 | Ru et al., 2014 |
| 125 | PL125 | Epiedesmin (ZINC03996196) | 7299790 | Ru et al., 2014 |
| 126 | PL126 | Valencene | 9855795 | Ru et al., 2014 |
| 127 | PL127 | (1S,5S)-1-isopropyl-4-methylenebicyclo[3.1.0]hexane | 11051711 | Ru et al., 2014 |
| 128 | PL128 | (5S)-5-[(1R)-1,5-dimethylhex-4-enyl]-2-methylcyclohexa-1,3-diene | 11127403 | Ru et al., 2014 |
| 129 | PL129 | (E)-5-(4-hydroxy-3-methoxy-phenyl)-1-piperidino-pent-2-en-1-one | 11630663 | Ru et al., 2014 |
| 130 | PL130 | (3R,8S,9S,10R,13R,14R,17R)-17-[(2R,5S)-5-ethyl-6-methylheptan-2-yl]-10,13-dimethyl-2,3,4,7,8,9,11,12,14,15,16,17-dodecahydro-1H-cyclopenta[a]phenanthren-3-ol (ZINC03982454) | 11870467 | Ru et al., 2014 |
| 131 | PL131 | Delta-elemene | 12309449 | Ru et al., 2014 |
| 132 | PL132 | (2R,4aR,8aR)-2-methyldecalin | 12816526 | Ru et al., 2014 |
| 133 | PL133 | (1R,5R,7S)-4,7-dimethyl-7-(4-methylpent-3-enyl)bicyclo[3.1.1]hept-3-ene | 13889654 | Ru et al., 2014 |
| 134 | PL134 | Calarene | 15560279 | Ru et al., 2014 |
| 135 | PL135 | Bisdemethoxycurcumin | 45934475 | Ru et al., 2014 |
| 136 | PL136 | 1,4-cadinadiene | 50986185 | Ru et al., 2014 |
| 137 | PL137 | Tricyclene | 55250308 | Ru et al., 2014 |
| 138 | PL138 | Alpha-Cubebene | 42608159 | Ru et al., 2014 |
| 139 | PL139 | Piperundecalidine | 44453654 | Ru et al., 2014 |
| 140 | PL140 | 3-phenylundecane | 20655 | Ru et al., 2014 |
| 141 | PL141 | 4-[(1-Carboxy-2-methylbutyl)amino]-2(1H)-pyrimidinone | 591989 | Ru et al., 2014 |
| 142 | PL142 | Bicyclo[3. 2. 2]non-6-en-3-one | N/A | Ru et al., 2014 |
| 143 | PL143 | Cedryl acetate | 9838172 | Ru et al., 2014 |
| 144 | PL144 | Isolongifolene epoxide | 107035 | Ru et al., 2014 |
| 145 | PL145 | N-isobutyleicosa-2(E),4(E),8(Z)-trienamide | N/A | Ru et al., 2014 |
| 146 | PL146 | Pisatin | 101689 | Ru et al., 2014 |
| 147 | PL147 | Tetradecahydro-1-methylphenanthrene | 609802 | Ru et al., 2014 |
| 148 | PL148 | Undulatone | 5281311 | Ru et al., 2014 |
| 149 | PL149 | Copaene | 25245021 | Ru et al., 2014 |
| 150 | PL150 | Linalool | 6549 | Ru et al., 2014 |
| 151 | PL151 | Sylvatine | N/A | Ru et al., 2014 |
| 152 | PL152 | beta-Cubebene | 93081 | Ru et al., 2014 |
| 153 | PL153 | (-)-Caryophyllene oxide | 1742210 | Ru et al., 2014 |
| 154 | PL154 | (-)-alpha-cedrene | 6431015 | Ru et al., 2014 |
| 155 | PL155 | (+)-Fargesin | CHEMBL462822 | http://www.genome.jp/db/pcidb |
| 156 | PL156 | Piperolactam A | CHEMBL387864 | http://www.genome.jp/db/pcidb |
| 157 | PL157 | (+)-Sesamin | CHEMBL252915 | http://www.genome.jp/db/pcidb |
| 158 | PL158 | 2-Phenylethanol | CHEMBL448500 | http://www.genome.jp/db/pcidb |
| 159 | PL159 | l-Zingiberene | CHEMBL479020 | http://www.genome.jp/db/pcidb |

### 3.2. Compounds classification and clustering

The result of the chemical classification shows that 159 phytochemicals were distributed among 26 different classes of compounds. There were 11 classes of compounds that contain only one phytochemical and these are carboxylic acids and derivatives (PL141), cinnamic acids and derivatives (PL68), isoflavonoids (PL146), napthalenes (PL72), organic nitrogen compounds (PL62), oxanes (Pl71), phenanthrenes and derivatives (PL147), phenol ethers (PL107), phenylpropanoic acids (PL69), pteridines and derivatives (PL67), pyridines and derivatives (PL17) and steroid and its derivatives (PL130). The adaptation of the plant against various abiotic and biotic stresses over the millions of years of evolution is responsible for such chemical diversity of the phytochemicals (Dias et al., 2012). Among all the classes, benzodioxole group constitutes the highest number of phytochemicals (31) and they are shown to be clustered together. It is the widely dispersed class of compounds among natural as well as synthetic drugs (Fang et al., 2011).

Hierarchical clustering of the phytochemicals is shown in Figure 1. The phylogenetic tree reveals that phytochemicals cluster with molecules that share similar scaffold. The class of chemical compounds corresponding to prenol lipids (condensation product of isoprene units) and fatty acyls are highly prominent in the dataset; a good agreement with the known fact that lipids form a large group of primary metabolites of plant (Busia, 2016).

**Figure 1.**
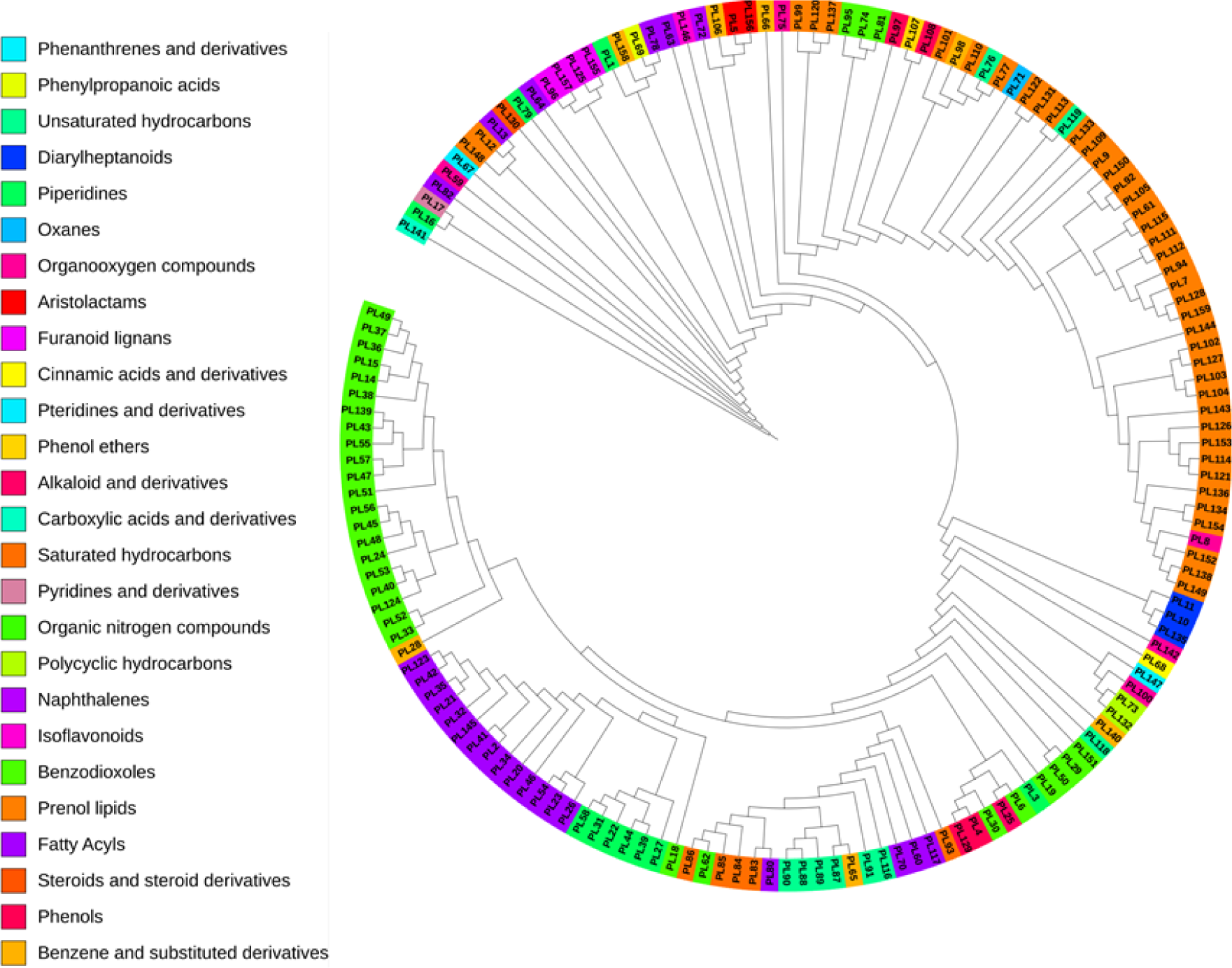
Hierarchical clustering of phytochemicals belonging to *Piper longum*. Phytochemicals were clustered on the basis of atom-pair descriptors and Tanimoto coefficient using Chemmine tool. It can be easily seen that most of the phytochemicals belonging to benzodioxoles, fatty acyl and prenol lipids category were clustered together.

### 3.2. Phytochemical – protein target (PC-PT) network

For understanding the interactions between small molecules and proteins, PC-PT bipartite network was constructed by mapping 159 phytochemicals to their potential proteins targets. This resulted in identification of 1109 unique human proteins that may be potentially targeted by the phytochemicals of *P. longum*. As may be seen in top two layers of tripartite network in Figure 2a and 3a, many phytochemicals are found to interact with multiple proteins, an effect known as polypharmacology.

**Figure 2.**
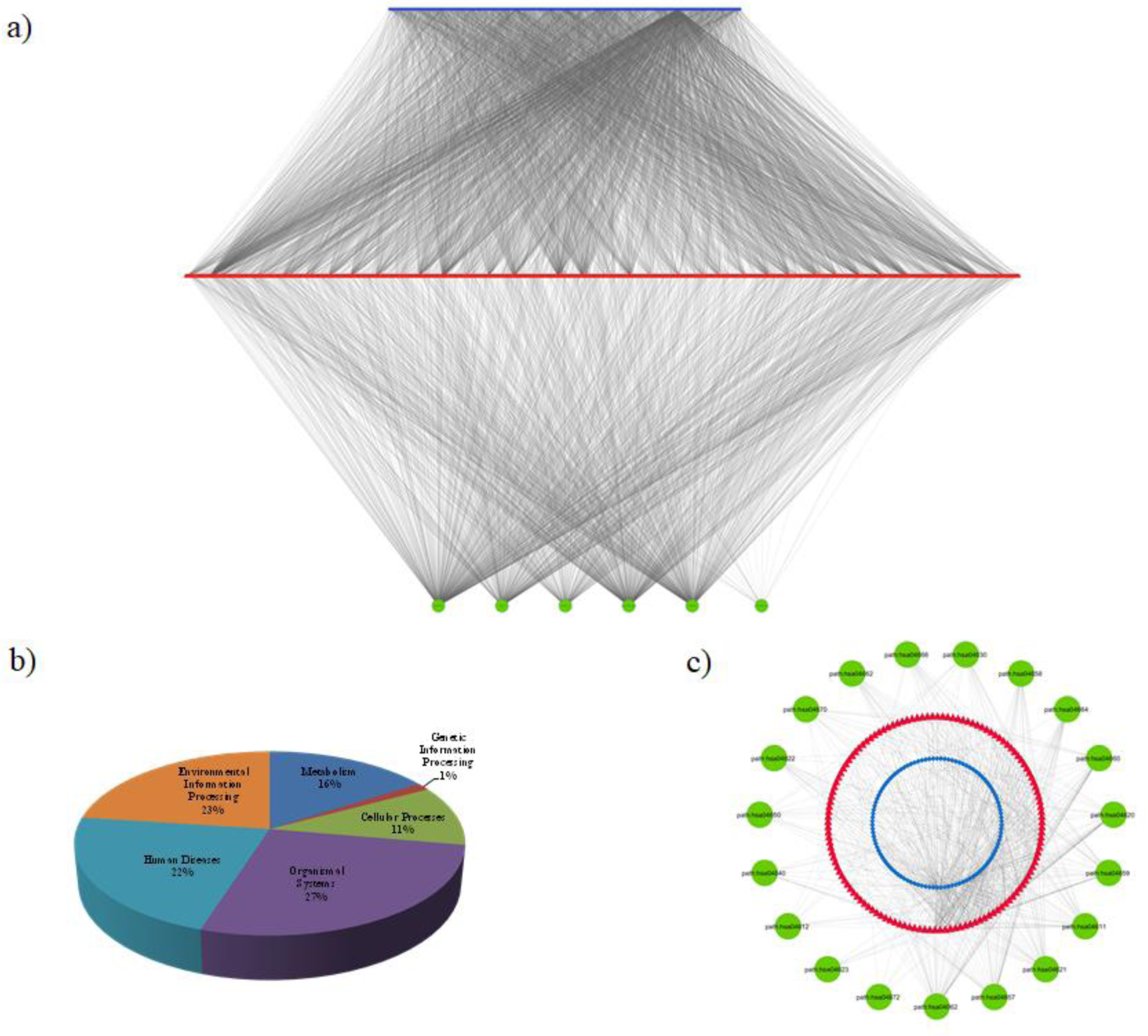
(a) A tripartite phytochemical - protein target - biochemical pathway (PC-PT-BP) network of the constituents of *Piper longum*. Top layer (blue) represents phytochemicals (159), middle layer (red) represents the potential protein targets (1109) and the third layer (green) represents association of protein targets with six pathway classes: metabolism, genetic information processing, environment information processing, cellular processes, organismal systems and human diseases. **(b) Involvement of target proteins of *Piper longum* in various human pathway classes.** Pathway mapping of target proteins were distributed among 6 pathway classes. Genetic information processing class includes pathways belonging to transcription, translation, replication & repair, folding, sorting and degradation. Environment information processing includes membrane transport, signal transduction, signaling molecules and their interactions. Cellular processes involve pathways of cell growth, cell death and transport (endocytosis, phagosome, lysosome etc.). Organismal system includes immune, endocrine, circulatory, digestive, excretory, nervous, sensory system. Human diseases and metabolism includes pathways associated with diseases and metabolic system (carbohydrates, lipids, amino acids etc.) respectively. **(c) Bioactives of *Piper logum* affecting the Human immune system**. The first layer (blue) represents 106 phytochemicals, regulating 11 immune pathways (green) by targeting 131 proteins (red).

**Figure 3.**
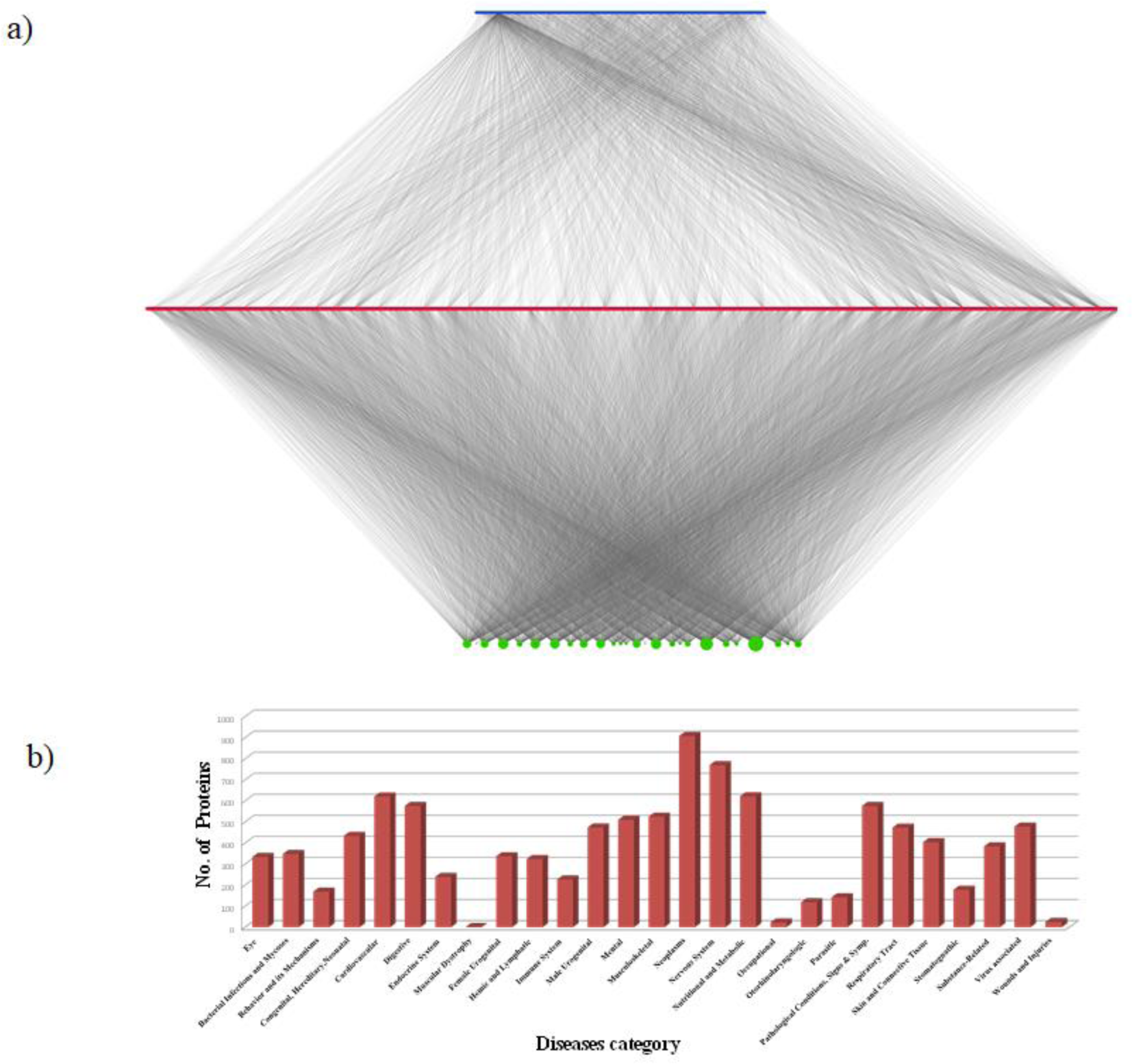
(a). Phytochemical - protein target - disease association (PC-PT-DA) network of *Piper longum*. Top layer (blue) represents phytochemicals **(159)**, middle layer (red) represents their protein targets **(1109)** and the bottom layer (green) represents association of protein targets with 27 disease classes obtained from DisGeNET. Size of the nodes in third layer (green) is proportional to their degree value. **(b) *Piper longum* protein targets and their involvement in 27 disease classes.** Number of potential protein targets associated with various disease classes as obtained from DisGeNETv4.0. Maximum numbers of targets are related with neoplasm and nervous system while least with the muscular dystrophy and occupational diseases.

Polypharmacological effect of the *P. longum* phytochemicals was evaluated using its PCt/Tt value (the average value of number of target for each compound). This value is calculated for each phytochemical identified, higher PCt/Tt value for a compound suggests that it may be activator or inhibitor of multiple proteins and may be individually or in combination serve as lead-compound. Numerous existing drugs are well known for their multi-targeting activities. One such example is Aspirin; which is usually used as an analgesic and at times also as an antipyretic (Vane et al., 2003), and its anti-inflammatory medication in treatment of various diseases like rheumatoid arthritis (Preston et al., 1989), pericarditis (Imazio et al., 2013) is also well known.

PL70 (palmitic acid) being ubiquitous in nature emerges out as the component with highest PCt/Tt value (0.55) in the network. Next in order phytochemicals are PL99, PL11, PL25 and PL66 with PCt/Tt value 0.068, 0.060, 0.057, 0,052 respectively. Network analysis shows that 99% of the phytochemicals are linked with more than 10 protein targets, indicating towards the multi-target properties of herb. However, sometimes the concept of polypharmacology could be a double-edged sword and may cause adverse effects when the idea is not fully understood (Reddy et al., 2013). In that case the approach may result in identifying new drug off-targets. Thus specificity of a compound to hit a unique target may be explored. Among 159 phytochemicals, PL59 (2, 2-Dimethoxybutane (C6H14O2)) is one such compound, that is shown to interact with single target (Uniprot ID - P08575). P08575 (receptor-type tyrosine-protein phosphatase C, CD45) plays a critical role in receptor-mediated signaling in both B and T-cells (Desai et al., 1990; McFarland et al., 1995). Since altered signaling process is one of the effect that leads to several disorders including SCID (Severe combined immunodeficiency syndrome). In such cases the explicit behavior of PL59 and its reliability to target specific protein can be appreciated.

Among 159 phytochemicals, 109 targets P10636. The high degree corresponding to this node measures its ability to interact with other phytochemicals. In other words, it is the most commonly targeted protein in the network. P10636 is a ―microtubule-associated protein tau (MAPT)‖ encoded by gene MAPT. A system level investigation of neurodegenerative dementia reveals the accumulation of the protein in diseased state, including frontotemporal dementia and supranuclear palsy (Caberlotto et al., 2014). Hence, it can be assumed that the role of identified 109 phytochemicals may contribute to the *P. longum* effect in the treatment of mental diseases. Although previous studies on treatment of mental disorders by the herbal extract is known (Hritcu et al., 2014) with special focus on piperine (PL25) and piperlongumine (PL68) in Parkinson‘s disease (Bi et al., 2015), detailed focus on the pharmacophore properties of additional phytochemicals identified will help in deciphering its detailed mechanism and the key assets that may be helpful in designing synthetic drugs. Additionally, the positive synergistic effect of the compounds can be explored for better affinity and efficacy.

Other proteins with high degree in the network include Q9NUW8, Q9NR56, Q5VZF2, Q9NUK0, P22303, and P06276 with value corresponding to 64, 60, 59, 57, 56 and 55 respectively. These proteins may receive special attention in aspects of behaving as key targets, specifically in light of the fact that interventions at specific protein can be weak in terms of binding affinity, yet they may be highly effective in combinations (Mason and J.S., 2012).

### 3.4. Phytochemical, Protein target and Biochemical pathway (PC-PT-BP) network

To obtain a global view of pathways targeted by *P. longum*, a tripartite network was constructed using its phytochemicals, their protein targets and associated biochemical pathways Figure 2a. 279 unique human pathway maps were classified into 6 broad categories i.e. metabolism, genetic information processing, environmental information processing, cellular processes, organismal systems and human diseases. The detailed mapping of target proteins into different pathways is given in Supplementary Table 1.

As suggested in Figure 2b, highest numbers of target proteins (455, 27%) are associated with pathways belonging to organismal systems, followed by environmental information processing category (384, 23%) and human diseases (371, 22%). This finding suggests that the mode of action of these phytochemicals may be largely *via* regulating organismal system (which includes immune, endocrine, circulatory, digestive, nervous and excretory system). Earlier *in vivo* study on *P. longum* also support this finding, in which it was shown that immunomodulatory properties of the herbal extract leads to increase in white blood cells (WBC) in Balb/c mice (Sunila and Kuttan, 2004). The endocrine effect is explained *via* the process of ovulation which include interrelationship between the endocrine and cytokine system. By modulating the inflammatory mediators like cytokines, reactive oxygen species etc., phytochemicals of this plant exert their antifertility properties (Lakshmi et al., 2006). Similarly, its effect on cardiovascular (Nabi et al., 2013), digestive (Ghoshal et al., 1996; Ghoshal and Lakshmi, 2002), nervous (Bi et al., 2015; Hritcu et al., 2014) and excretory (Meena et al., 2009) system have also been studied. Protein targets such as P28482, Q9Y243, P31751, P31749, P42336, P27986 and P27361 are found to be involved in many pathways. Thus it may be hypothesised that these proteins may be important target as their modulation may lead to regulation of multiple pathways.

To explore the basic principle of the herb in relation to the immune system of the human, a sub network of immune pathways being regulated by *P. longum* was created. 106 phytochemicals of 159 are shown to regulate 19 immune pathways of human *via* 131 proteins. 57.25% of these proteins are affecting the immune system *via* chemokine (hsa04062) and interleukin (hsa04657) signaling pathways. Role of piperine in reducing Th2 cytokines and regulating cytokine in asthma models have been explored earlier (SeungHyung and YoungCheol, 2009).

This sub-network identifies Prostaglandin G/H synthases (PTGS) as common targets. PTGS-1 (P23219) and PTGS-2 (P35354) are the targets of 28 and 22 phytochemicals respectively. This enzyme is involved in conversion of arachidonate to prostaglandin H2, which are the major components that induce inflammation and pain (Ricciotti and FitzGerald, 2011). The high rate of edema inhibition by herb-oil in comparison to the standard anti-inflammatory drug, Ibuprofen is also reported (Kumar et al., 2009).

Further, the network data provides that NOD-like receptors and Toll-like receptors (TLRs) are regulated through 33 and 29 targets respectively. The receptors play a key role in innate immunity. The pathogen invasion caused by bacterial lipopolysaacride (LPS) induces signalling pathways which further lead to the activation of macrophages *via* TLRs. Since an earlier report on the herb shows that the root area possesses anti-amoebic properties (Ghoshal and Lakshmi, 2002). Thus, the sequence of events which lead to interaction of receptor proteins with these phytochemicals (especially present in root region) can be focussed to explore their effect on the innate immunity.

Although, various studies have supported the anti-inflammatory behaviour of *P. longum*, our work reveals that this effect is not due to a limited number phytochemicals rather a vast number of phytochemicals are involved in this property. Among 105 phytochemicals involved in immunomodulation, palmitic acid (PL70), demethoxycurcumin (PL11), bisdemethoxycurcumin (PL10), 1,2-benzenedicarboxylic acid (PL66), sesamin (PL96) and piperine (PL25) are the top-immunomodulators with 76, 15,12, 11, 11, 10 protein targets respectively. Thus, combining the effect of other phytochemicals reported in our study will help to provide a whole-sum picture of the herb‘s immunomodulatory potency. Additionally, the use of analytical and structural chemistry of the phytochemicals and phytochemical-protein target complexes will help in understanding the molecular mechanism in detail. The detailed information of the immune pathways considered and the number of target proteins involved in each class is represented in Table 2.

**Table 2.**
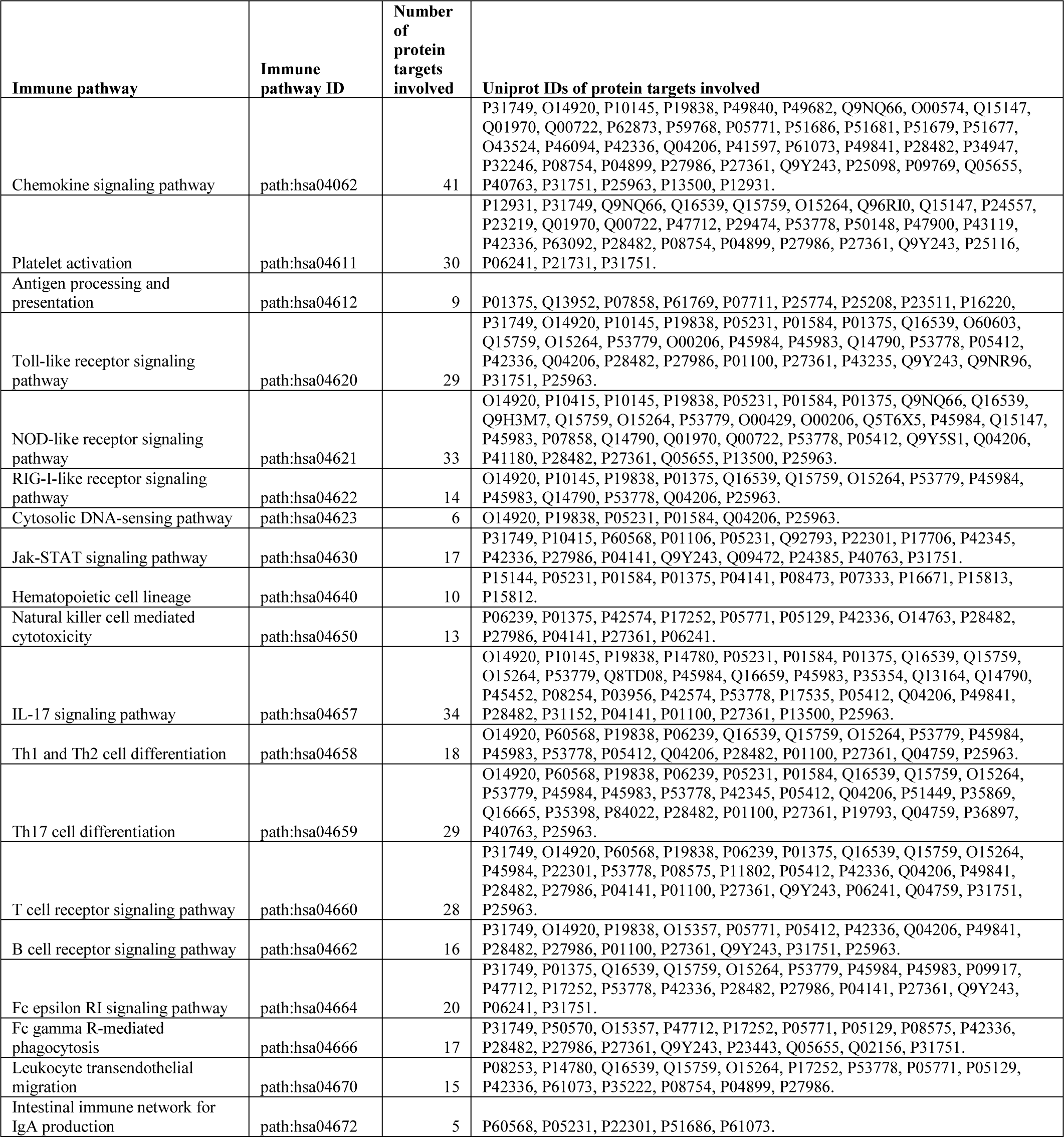
List of protein targets of *P. longum's* biochemicals involved in immune pathways of *Homo sapiens*.

### 3.5. Phytochemical, Protein target and Disease association (PC-PT-DA) network

The data of protein targets and the diseases in which they are involved was collected and a tripartite network was drawn (Figure 3a). The diseases were classified into 27 classes and distribution of proteins among each class is shown in Figure 3b. It can be easily seen that majority of protein targets have their association with neoplasm and nervous system diseases, with 908 and 770 proteins respectively. Numerous studies have shown the effect of pepper plant in treatment of different cancers including prostate, breast, lung, colon, lymphoma, leukemia, primary brain tumors, and gastric cancer. A recent study states that the plant‘s anti-cancerous property is due to the inhibitory mechanism of Glutathione S-transferase pi 1 (GSTP1) by its compound piperlongumine (PL29). GSTP1 is overexpressing protein in cancerous cells and the reactive olefins in the piperine (PL25) attenuate the cancer-cell proliferation by blocking its active site (Harshbarge et al., 2017).

According to the network data, association between *P. longum* and nervous system diseases is mainly by the regulation of 434 protein targets. Among these, majority of proteins are involved in signaling transmission and developmental pathways. Out of total 434, 29 protein targets are found to be interacting with piperine (PL25) that is known to have antiepileptic (Pattanaik et al., 2006), analgesic, anticonvulsant (Bukhari et al., 2013) nature. Anti-depression like activity of this compound on animal samples suggests that the compound may act a potential functional food to improve brain function (Wattanathorn et al., 2008).

Analysis of degree distribution shows that 1,2-Benzenedicarboxylic acid (PL66), demethoxycurcumin (PL11) and D-camphor (PL99) are having more number of protein interactors in comparison to piperine (PL25) i.e 36, 34 and 33 respectively. Thus, it may be inferred that these phytochemicals may also have effect in neuroprotection and may act as new lead compounds for neurodegenerative disorders.

The prioritization of the target proteins of *P. longum* was performed by mapping them to the approved drug targets of DrugBank with intent of selecting known drug targets for identifying their novel regulators from this herb‘s constituents. For this, common target proteins among all the four target prediction softwares were selected. Five proteins (P04150, P37231, Q8NER1, P21397 and P27338) among these are FDA approved drug-targets, reflecting their importance for identification of their novel regulators. The position of these potential targets in the PC-PT-DA network shows that, 77 phytochemicals are involved in their regulation (Figure 4).

**Figure 4.**
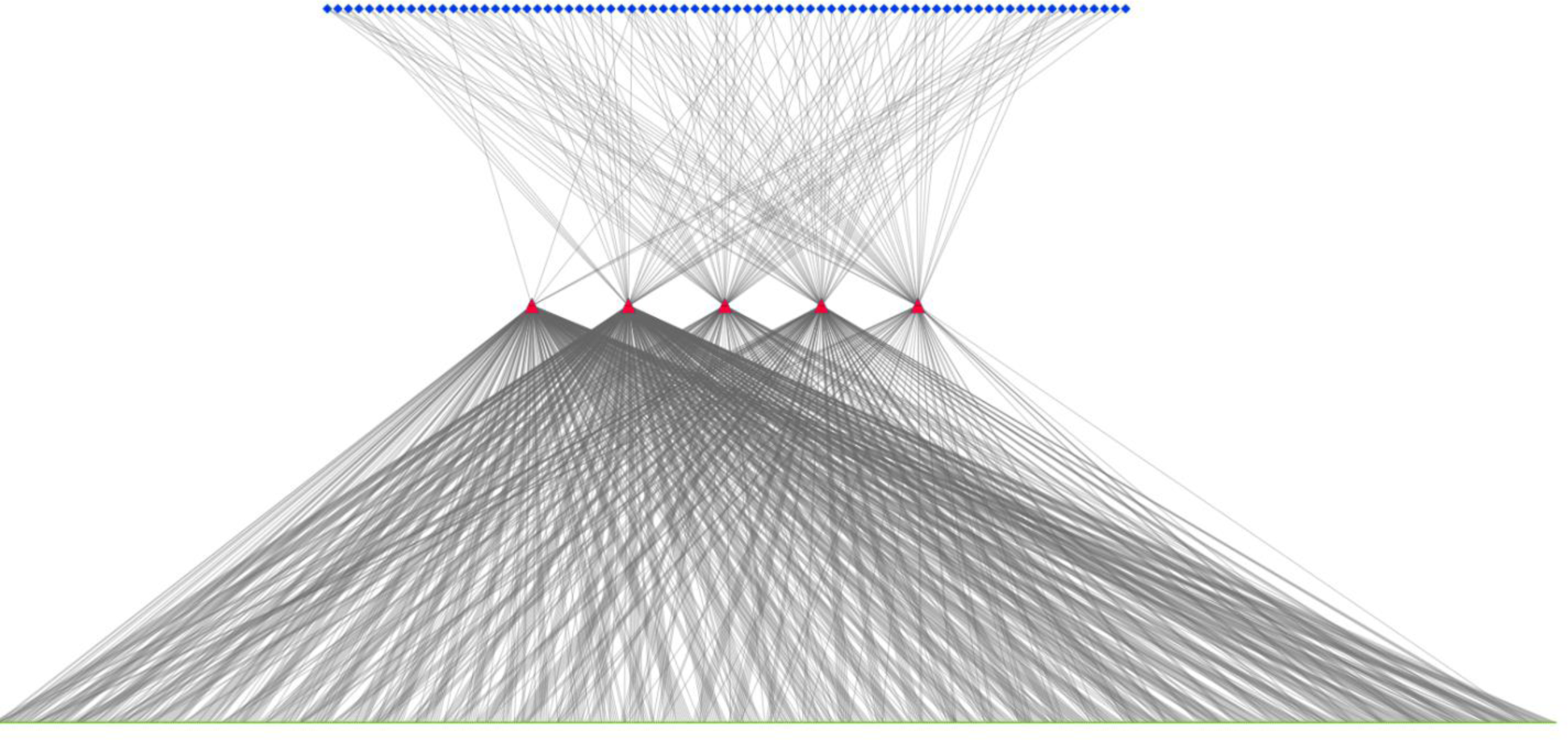
Potential FDA-approved targets from target dataset of *Piper longum*. Middle layer (red) represents the 5 key protein targets, which are regulated by 77 phytochemicals of *Piper longum,* represented in the top layer (blue). Third layer (green) represents the mapping of these targets in 825 diseases.

Further, these key targets are associated with diverse array of diseases that means they are involved in multiple levels in the biological system. 28 protein targets were predicted by three of the four softwares used for phytochemical-protein interaction prediction. These may be considered as novel targets as compared to DrugBank entries and may be further explored with respect to the phytochemicals involved in their regulation (Supplementary table 3). The network data provides a scope to explore the interrelationship of the target proteins with phytochemicals using computer aided drug discovery approaches, which will be helpful in understanding the medicinal and multi-targeting potential of the herb in detail. Further, pleiotropic nature of the genes at the system level can be investigated.

### 3.6. Module and GO enrichment analysis of human PIN

First degree interactors of all the target proteins of *P. longum* were used to construct a subnetwork of PIN of *Homo sapiens* (Figure 5). Topological analysis of the network shows that its degree distribution follows a power-law with^−1.245^. The PIN was analysed using MCL clustering algorithm for module identification. Modules are shown in supplementary figure 1. Functional enrichment analysis of the modules with dense connection shows that *P. longum* exerts its effect mainly through regulating cell cycle, signal transduction, genetic information processing and metabolism machinery (Table-3)

**Figure 5.**
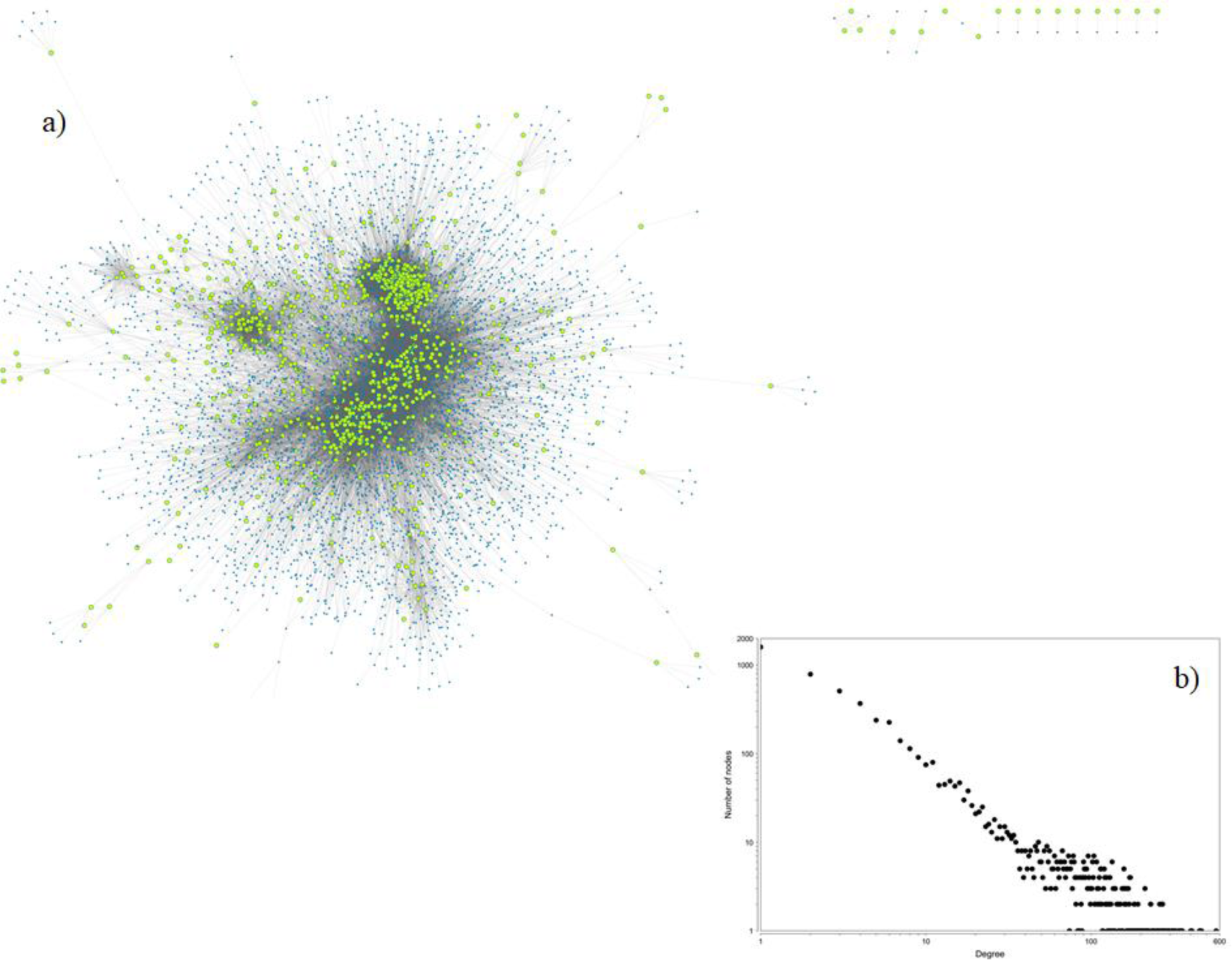
(a) Protein-protein interaction subnetwork of *Homo sapiens* targeted by phytochemicals of *P. longum*. First neighbours of all the targets proteins were mapped into the human PPI as obtained from STRING having high confidence level (score ≥ 900). Green highlighted nodes in the network represent the location of the target proteins of *P. longum.* **(b) Node degree distribution of the PPI subnetwork.** The neighbourhood connectivity of each node is represented using node degree distribution graph, analysed using Cytoscape. Both axes of the graph are represented in the logarithmic scale.

**Table 3.** GO based biological processes of 15 highly connected modules in the protein-protein interaction subnetwork of *Homo sapiens* targeted by phytochemicals of *P. longum*.

| Cluster-Number | GO-ID | P-value | Description |
| --- | --- | --- | --- |
| cluster-1 | 42770 | 1.61E-29 | DNA damage response, signal transduction |
| cluster-2 | 23052 | 7.10E-28 | Signaling |
| cluster-3 | 278 | 9.18E-122 | Mitotic cell cycle |
| cluster-4 | 7186 | 2.09E-122 | G-protein coupled receptor protein signaling pathway |
| cluster-5 | 6325 | 8.92E-63 | Chromatin organization |
| cluster-6 | 7166 | 2.42E-47 | Cell surface receptor linked signaling pathway |
| cluster-7 | 22900 | 2.05E-118 | Electron transport chain |
| cluster-8 | 16055 | 1.77E-27 | Wnt receptor signaling pathway |
| cluster-9 | 19220 | 1.48E-14 | Regulation of phosphate metabolic process |
| cluster-10 | 6357 | 7.67E-21 | Regulation of transcription from RNA polymerase II promoter |
| cluster-11 | 6096 | 8.75E-33 | Glycolysis |
| cluster-12 | 6260 | 2.33E-48 | DNA replication |
| cluster-13 | 9889 | 2.84E-14 | Regulation of biosynthetic process |
| cluster-14 | 3735 | 1.28E-127 | Structural constituent of ribosome |
| cluster-15 | 7049 | 1.44E-41 | Cell cycle |

Replication and repair mechanism (Module-1) involve proteins such as ATM, BLM and BRCA2. ATM is serine/threonine kinase gene and acts as an important cell-cycle check point kinase (Abraham, 2001). BLM gene is a Bloom syndrome RecQ like helicase; protein encoded by this gene is involved in suppression of inappropriate recombination event in the cell (Hickson, 2003). BRCA gene helps in maintaining the stability of the genome. BRCA2 is involved in double stranded DNA repair by regulating the homologous recombination pathway (Karran, 2000). Mitotic cell cycle (Module-3) contains gene like ANAPC10, CDC 20 etc. ANAPC10 is a core subunit of anaphase promoting complex and it is known that APC genes get altered in human colon cancer (Wang et al., 2003). CDC20 (cell division cycle 20) acts as a regulatory protein and it has been shown *in vitro* that it is a promising therapeutic target for cancer treatment (Wang et al., 2013). This shows that *P. longum* anti-cancerous activities are mainly due to regulation of cell-cycle events and DNA-repair mechanisms.

It is commonly observed that disease occurrence is associated with the signal transduction failure, but the degree of its association varies greatly depending on the severity of disease. Wnt signalling pathways (Module-8) contain genes such as APC2, WNT7A and FRAT-1. Mutation in Wnt signalling pathway genes like APC (Adenomatous polyposis coli) are particularly evident in memory deficit cases.

Gene knockout studies on the mice sample shows that the gene plays an important role in the regulation of spinal locomotor activity and memory performance (Koshimizu et al., 2011). Its involvement in ocular, bone density disorders (Canalis, 2013; MacDonald et al., 2009) and colorectal cancer (Schneikert and Behrens, 2007) are also well studied. The proteins of WNT gene family like WNT7A (Wnt family member 7A) encode secreted signalling proteins (Smolich et al., 1993). An earlier report in the developmental biology shows that Wnt7a is a highly conserved gene and play an important role in early development of midbrain and telencephalon region of human brain (Fougerousse et al., 2000). Target data analysis shows that ―P12644‖ encoded by BMP4 (Bone morphogenetic protein-4) is the target gene of WNT/beta-catenin signalling pathway (Kim et al., 2002). Thus, it may be hypothesised that the herb‘s nootropic effect is mainly associated with the proteins constituting this module.

*P. longum*‘s anti-inflammatory quality may be linked to G-protein coupled signalling process (Module-4). This module contains genes like GRK2 and GRK6. Inflammatory mediators modulate GRKs signaling either by transcription regulation or its degradation. GRK2 act as mediator in the pathway that causes inflammatory pain (Sun and Richard, 2012). Thus, targeting GRKS by *P. longum* could be the reason of its anti-inflammatory properties. Pain relieving properties of this herb can be easily linked to the proteins in Module-4.

Proteins of module 10 and 14 are involved in genetic information processing by regulating transcription and translation machinery respectively. Genes like HES1, RUNX1, TGFB1, EP300 constitute module 10. HES1 is a hes family bHLH transcription factor-1. RUNX-1 is runt related transcription factor 1 which participates in haematopoiesis. Its involvement in leukaemia conditions is well documented (Huang et al., 2009). TGFB1 is a transformation growth factor beta-1 and it regulates cell-proliferation, differentiation and unregulated in tumor cells (Cohen, 2003). EP300 encodes E1-A associated cellular p300 transcriptional co-activator protein. It also helps in stimulating hypoxia-induce genes. A defect in the gene leads to Rubinstein-Taybi syndrome (Roelfsema et al., 2005). Thus, *P. longum* may be effective in improving transcriptional errors or the diseases associated with it.

Module 14 constitutes ribosomal proteins like RPL36, RPS3, RPL8, RPS7, RPS13 and RPL3. These proteins help in maintaining the structural integrity of ribosomal assembly. Extraribosomal function of ribosomal proteins include regulation of gene expression, cell-cycle control, regulation of apoptosis, modulation of DNA repair, regulation of development and differentiation, modulation of cell migration and invasion and regulation of angiogenesis (Wang et al., 2015). Dysfunction of ribosome leads to a condition known as ribosomopathies. Although no data supports the herb‘s effect in treating such dysfunction, but we believe that this area needs a detailed investigation. *P. longum* phytochemicals may have a regulatory effect on ribosomopathies.

Module-7 proteins are present in the metabolic machinery which is reported to regulate Electron transport chain (ETC) and forms basis of the energy production in the cell. These proteins are MT-CO1, UQRC1 and NDUFA6. MT-CO1 (mitochondrial cytochrome c oxidase subunit-1), UQRC1 (ubiquonol-cytochrome c reductase core protein) and NDUFA6 (NADH ubiquinone reductase subunit A6) are associated with the redox processes of ETC (Herrero–Martín et al., 2008; Cadenas et al., 1977; Zickermann et al., 2014) and aid in an essential aspect of ATP generation. Abnormalities in ETC chain are characteristic feature of Alzheimer‘s (Parker et al., 1994) and Parkinson‘s disease (Parker et al., 1989) of brain. Thus, the chemical compounds produced in *P. longum* are involved in the crucial steps of ETC.

Glucose metabolism (Module-11) includes enzymes that are crucial for the conversion of glucose into pyruvate. This process is carried out by the cell to meet its energy requirements. The module contain genes like PGK2 (Phosphoglycerate kinase 2), ENO1 (Enolase 1), PKLR (Pyruvate kinase) and ALDOA (Aldolase, fructose-bisphosphate A). This shows that the herb may also be helpful in regulating energy metabolism. Additionally, high rate of glycolysis have been reported in cancerous cells (Elstrom et al., 2004). In such condition *P. longum* anti-cancerous property may be correlated with its interaction with modules involved in glucose metabolism.

### 3.7. Drugability analysis and docking studies

The estimation of ADMET and other drug like properties are important to consider at early stage of drug-discovery process as majority of the drug candidates fail the clinical trial due to poor ADMET properties (Van De Waterbeemd and Gifford et al., 2003). We evaluated the complete pharmacokinetic and toxicity profile of each phytochemical of *P. longum* for characterizing their drug likeliness (Supplementary Table 2).

*In silico* estimation shows that the percentage intestinal absorption of the phytochemicals is more than 90% for 136 phytochemicals. 93% of the phytochemicals are likely to permeate the Caco2 cells as they have permeability value greater than 1. Longumosides B (PL13), piperazine adipate (PL63), 2-amino-4-hydroxypteridine-6-carboxylic acid (PL67) and 4-[(1-Carboxy-2-methylbutyl) amino]-2(1H)-pyrimidinone (PL141) are predicted to have least permeability among other phytochemicals. The possible distribution of compounds through various compartments of the body was accessed using its Blood-brain barrier (BBB) penetration and Central nervous system (CNS) penetration coefficient. One of the criteria for a successful CNS drug is that the compound should not have binding affinity for P-glycoprotein (Pajouhesh and Lenz et al., 2005). The result shows that 111 phytochemicals may act as non-substrate of P-glycoprotein, thus possess the ability to be a potential CNS drug. Piperine (PL25) is known to have CNS acting power (Lee et al., 2005), the BBB and CNS penetration values were found to be −0.131 and −1.932 respectively. 115 phytochemicals of *P. longum* were showing the BBB and CNS penetration values similar to or greater than the piperine (PL25) values. This indicates that CNS targeting potential of other phytochemicals is noteworthy and can be explored further.

**Figure 6.**
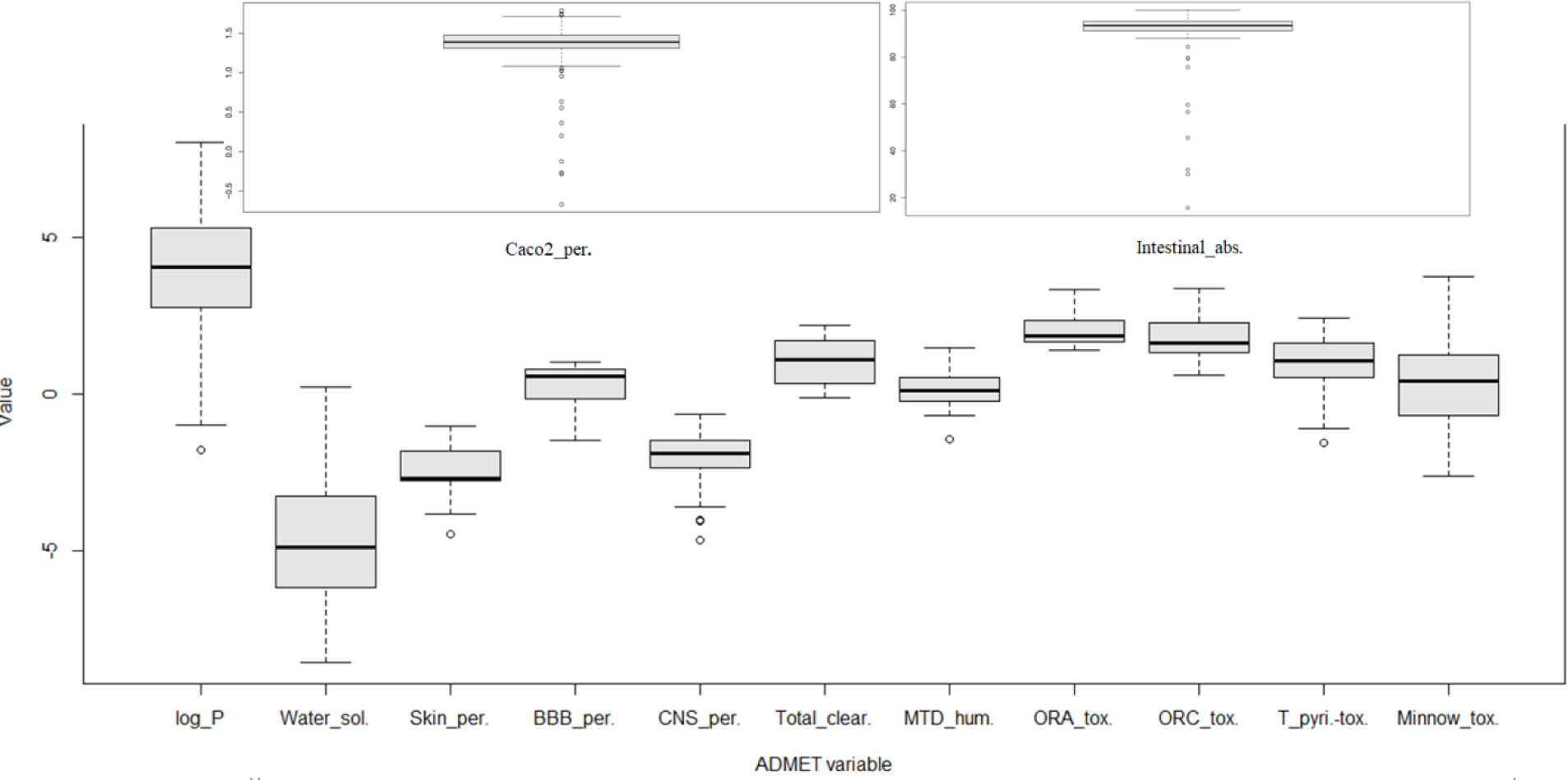
Box and Whisker plots showing ADMET properties distribution of the *Piper longum's* phytochemicals. The graph shows the plots of ADMET variables corresponding to octanal-water partition coefficient (log_P), water solubility (Water_sol.), skin permeability (Skin_perm.), blood-brain permeability (BBB_per), central nervous system permeability (CNS_perm.), total clearance (Total_clear.), maximum tolerated dose-humans (MTD_humans), oral rat acute toxicity (ORA_tox.), oral rat chronic toxicity (ORC_tox.), *T. Pyriformis* toxicity (T_pyri. tox), minnow toxicity (Minnow_tox), caco2 cell permeability (Caco2_per.) and intestinal absorption (Intestinal_abs.).

Lipinski‘s ―rule of five‖ criterion was adopted to estimate the likeliness of the phytochemicals to act as drug molecule (Lipinski et al., 1997). 105 phytochemicals are shown to maintain the criteria of molecular weight less than 500 Dalton, number of hydrogen bond donors less than 5, hydrogen bond acceptors less than 10 and logP value (octanal-water partition coefficient) less than 5. Toxicity risk was also evaluated by checking Ames toxicity, oral-rat acute toxicity, oral-rat chronic toxicity, hepatotoxicity, cardiotoxicity, *T. pyriformis* toxicity and Minnow toxicity. 13% of the compounds are predicted to be positive for hepatotoxicity. The cardiotoxicity was evaluated with hERG (human ether-a-go-go related gene) inhibition. Interestingly, not a single phytochemical is positive for hERG1 inhibition, which reflects the cardio-protective nature of *P. longum*.

Final screening of all the parameters in phytochemical dataset resulted in identification of 20 phytochemicals that possess high probability of acting as effective lead molecules (Supplementary Table 2). For identifying the novel regulatory molecules to the previously discussed 5 key protein targets, these 20 phytochemicals were back-mapped in the PC-PT network. This resulted in selection of 7 phytochemicals that were forming 16 phytochemical-protein target pairs. To estimate the binding affinity of these potential phytochemical-protein target pairs, docking studies were performed. This is essential to identify the best fit between the phytochemical and protein molecule, both in terms of energy and geometry. Binding energy calculations of each pair is represented in Table 4. Chemical features of all these phytochemicals can be explored further to design their synthetic analogues with optimised pharmacological activity.

**Table 4.** Docking energies of potential protein targets with their corresponding regulatory phytochemicals.

| Protein Targets (Uniprot ID) | PDB ID | Phytochemical ID | Docking energy (Kcal/mol) |
| --- | --- | --- | --- |
| P04150 | 4MDD | PL125 | -4.72 |
| P21397 | 2BXR | PL6 | -8.94 |
|  |  | PL61 | -4.68 |
|  |  | PL81 | -4.49 |
|  |  | PL95 | -4.39 |
|  |  | PL125 | -8.47 |
|  |  | PL152 | -7.4 |
| P27338 | 2BXS | PL6 | -8.71 |
|  |  | PL61 | -4.31 |
|  |  | PL81 | -4.84 |
|  |  | PL95 | -4.63 |
|  |  | PL125 | -7.7 |
|  |  | PL152 | -7.19 |
| P37231 | 4EMA | PL125 | -7.6 |
| Q8NER1 | 3J5P | PL6 | -9.04 |
|  |  | PL71 | -4.59 |

## 4. Summary

Traditional Indian medicine (TIM) system commonly known as Ayurveda is about 5000 BC old heritage of Indian subcontinent and is a huge repository of information about multiple natural medicines for their therapeutic potential. *P. longum* is an important constituent of many Ayurvedic formulations; the most widely used among these is ―Trikatu‖. However, the multi-targeting potential of this herb and underlying mechanism of its cellular-level action are still unexplored. The approach of network pharmacology in recent years has provided a way to investigate the healing potential of the traditional herbs for drug discovery and drug development procedures. Therefore, in the current study, we have examined the medicinal effect of *P. longum* using network pharmacology. The methodology involves literature survey, database mining, drug-likeliness prediction, phytochemical-protein target and protein target-disease relationship study to examine the multifaceted potential of this herb. The extensive text mining of its phytochemicals and their effect on the human proteins (involved in diseases) has led to the identification of novel phytochemical-protein target relationships. Our study highlights 5 key FDA approved targets and their 77 regulating phytochemical partners, for which chemical scaffold analysis can be performed to investigate the important drug-ligand interactions at the atomic level. Immune pathway network analysis has led to the identification of top immunomodulatory phytochemicals which involves palmitic acid (PL70), demethoxycurcumin (PL11), bisdemethoxycurcumin (PL10), 1, 2-benzenedicarboxylic acid (PL66), sesamin (PL96) and piperine (PL25).

Further, module analysis in the human protein-protein interaction network was carried out to decipher the effects of phytochemicals at global perspective. Significant modules include proteins involved in the regulation of cell cycle, signal transduction, metabolism and genetic information processing. Among 159 phytochemicals, 20 are estimated to be effective lead molecules based on drug-likeliness filter. We believe that our work based on systems level network assisted studies of *P. longum* will offer a new way to look upon the hidden potentials of this herb. The data obtained from docking analysis may be taken for *in vitro* studies, which may eventually be helpful in identification of novel and effective lead molecules from natural pool of compounds present in *P. longum*. We are hopeful that this study will prove to be an important basis for understanding the phytochemical-protein level coordination in various Ayurvedic formulae that uses *P. longum* as an integral part. Further, the phytochemical-protein target and disease relationship represented in the form of interaction networks will be helpful in understanding the underlying molecular mechanism in detail.

## Acknowledgements

N.C. is grateful to the University Grant Commission, India for providing PhD fellowship and V.S. thanks Central University of Himachal Pradesh for providing computational infrastructure.

## Author's contributions

V.S. conceptualized and designed this study. N.C. performed the computational analysis. Both the authors analyzed the results and wrote the manuscript.

